# Iota-carrageenan and Xylitol inhibit SARS-CoV-2 in cell culture

**DOI:** 10.1101/2020.08.19.225854

**Authors:** Shruti Bansal, Colleen B. Jonsson, Shannon L. Taylor, Juan Manuel Figueroa, Andrea Vanesa Dugour, Carlos Palacios, Julio César Vega

## Abstract

COVID-19 (coronavirus disease 2019) is a pandemic caused by SARS-CoV-2 (severe acute respiratory syndrome-coronavirus 2) infection affecting millions of persons around the world. There is an urgent unmet need to provide an easy-to-produce, affordable medicine to prevent transmission and provide early treatment for this disease. The nasal cavity and the rhinopharynx are the sites of initial replication of SARS-CoV-2. Therefore, a nasal spray may be a suitable dosage form for this purpose. The main objective of our study was to test the antiviral action of three candidate nasal spray formulations against SARS-CoV-2. We have found that iota-carrageenan in concentrations as low as 6 µg/ mL inhibits SARS-CoV-2 infection in Vero cell cultures. The concentrations found to be active in vitro against SARS-CoV-2 may be easily achieved by the application of nasal sprays already marketed in several countries. Xylitol at a concentration of 5 % m/V has proved to be viricidal on its own and the association with iota-carrageenan may be beneficial, as well.

## Introduction

SARS-CoV-2 is a single stranded positive sense RNA virus responsible for COVID-19. COVID-19 has become one of the worst pandemics of our time counting more than 12.232,700 confirmed cases and more than 554,000 deaths worldwide by July 10^th^, 2020 [1]. In most cases, COVID-19 manifests itself clinically with flu-like symptoms as a mild or uncomplicated illness, eventually resolving spontaneously. However, 15% of patients develop severe pneumonia that requires hospitalization and oxygen support, and 5% of them need admission to an intensive care unit (ICU). More than half of this patients may die [2]. Even children can be affected although with milder symptoms than adults and can be transmitters of the disease. [3]. There are no adequate therapeutic or preventive medicines available, so effective therapeutic approaches are urgently needed to reduce the spread of the virus and its death toll.

During the first days of disease the virus is localized mainly in the nasal cavity and the nasopharynx [4,5]. Recent data show that high viral load and a long virus-shedding period was associated with severe COVID-19 [6, 7]. Therefore, the use of antiviral nasal sprays would contribute to reduce nasal and nasopharyngeal viral load, thus slowing down the disease progression in the treated patient and the disease transmission to others in close contact with him or her.

Carrageenans are linear sulfated polysaccharides that are often extracted from red seaweeds. Carrageenans are commercially available in the form of kappa (κ), iota (ι) or lambda (λ). They have been used for years as thickening agents and stabilizers for food. At present they are extensively used in the food (cold cuts, cheese, etc.) and in the cosmetic and pharmaceutical industry as suspension and emulsion stabilizers. Their antiviral capacity has been described decades ago and has been experimentally confirmed on herpes virus type 1 and 2, human papiloma virus, H1N1 influenza virus, dengue virus, rhinovirus, hepatitis A virus, enteroviruses, and coronaviruses. Iota-carrageenan inhibits several viruses based on its interaction with the surface of viral particles, thus preventing them from entering cells and trapping the viral particles released from the infected cells. [8, 9, 10, 11, 12, 13].

Iota-carrageenan formulated into a nasal spray has proved to be safe and effective against virus causing common cold [14, 15, 16]. In vitro studies in cell cultures (HeLa and Calu-3) and in primary respiratory epithelial cells have shown inhibition of rhinovirus, Influenza virus and common-cold coronavirus. Iota-carrageenan is most active against common-cold coronavirus, inhibiting the infection up to 90 %. An iota-carrageenan spray reduced mortality by at least 50 % in mice infected with lethal doses of H1N1 influenza virus [17]. In all cases the antiviral action of iota-carrageenan is more effective, when administrated preventively or in the early stages of disease and has shown synergy with other antiviral agents. Studies performed on adults and children with common cold demonstrate effectivity of iota-carrageenan nasal spray to alleviate the clinical symptoms and shorten their duration, as well as to decrease the viral load of nasopharyngeal specimens and the relapses during the follow-up period [14, 15, 16, 18, 19, 20]. Iota-carrageenan-containing nasal sprays are already on the market in several countries in the world.

Xylitol is a polyol that has been used as a sugar substitute in Finland since the 1960s. It is a polyol, (formula CHOH)3(CH2OH)2), which is obtained from xylan extracted from hardwood, which has demonstrated multiple health benefits [21]. It has been extensively used in buccal health care to prevent caries because of its antibacterial capacity. It is already being used in otorhinolaryngology as a nasal spray and lavage for the treatment of rhinosinusitis and the prevention of otitis media. [22,23]. Studies “in vitro” and in animal models has shown antiviral properties of Xylitol against human respiratory syncytial virus (24).

Both iota-carrageenan and xylitol are safe for humans, being used in much larger amounts as food additive and sweetener, respectively, than those that may be used for nasal delivery. Nasal safety of iota-carrageenan by nasal and nebulization administration has been already confirmed empirically [25]. The same holds for 5 % Xylitol water solution both applied as nasal spray and nasal irrigation [26], as well as applied as a nebulization solution [27]. Both are included in nasal formulations already on the market for use in children and adults.

Based on the above knowledge, an experiment was designed and carried out in a Biosafety Level 3 (BSL3) laboratory to investigate the SARS-CoV-2 inhibition capacity of three different candidate preservative-free nasal formulations.

## Materials and methods

### Cells and Virus

Vero E6 cells were purchased from the American Type Culture Collection. Vero E6 cells were grown in complete minimal essential media (c-MEM) (Corning, NY, USA) which included 5% fetal bovine serum (FBS)(Gibco, Waltham, MA, USA), 5 mM penicillin/streptomycin (Gibco), and L-glutamine (Gibco). Cells were incubated at 37°C with 5% CO2. SARS-CoV-2 Isolate USA-WA1/2020, was obtained from BEI Resources (catalogue number NR-52281, Manassas, VA, USA). Virus master seed stock was prepared in T175 flasks of Vero E6 cells using a multiplicity of infection (MOI) of 0.1. Each flask was harvested on day two post-infection and supernatant was centrifuged twice at 220 × g for 15 minutes to remove cellular debris. Titer of virus stock was determined by plaque assay on Vero E6 cells.

### Preparation of sample formulations

All the formulations and placebos were prepared at Laboratorio Pablo Cassará S.R.L. (Argentina) under aseptic conditions and provided by Amcyte (US) to the University of Tennessee Health Science Center. Composition of different formulations is depicted in Tables 1 and 2. Samples 1, 2 and 3 were diluted at stock concentrations of 1200 µg/mL, 120 µg/mL, 12 µg/mL and 1.2 µg/mL using samples P1, P2 and P3 respectively as diluents. To determine antiviral efficacy of formulations by titer reduction assay, sample formulations were used at a final iota-carrageenan concentration of 600 µg/mL; 60 µg/mL, 6 µg/mL and 0.6 µg/mL. Equivalent concentration of placebos (samples P1, P2 and P3) was used for titer reduction assay as controls.

**Table 1.**
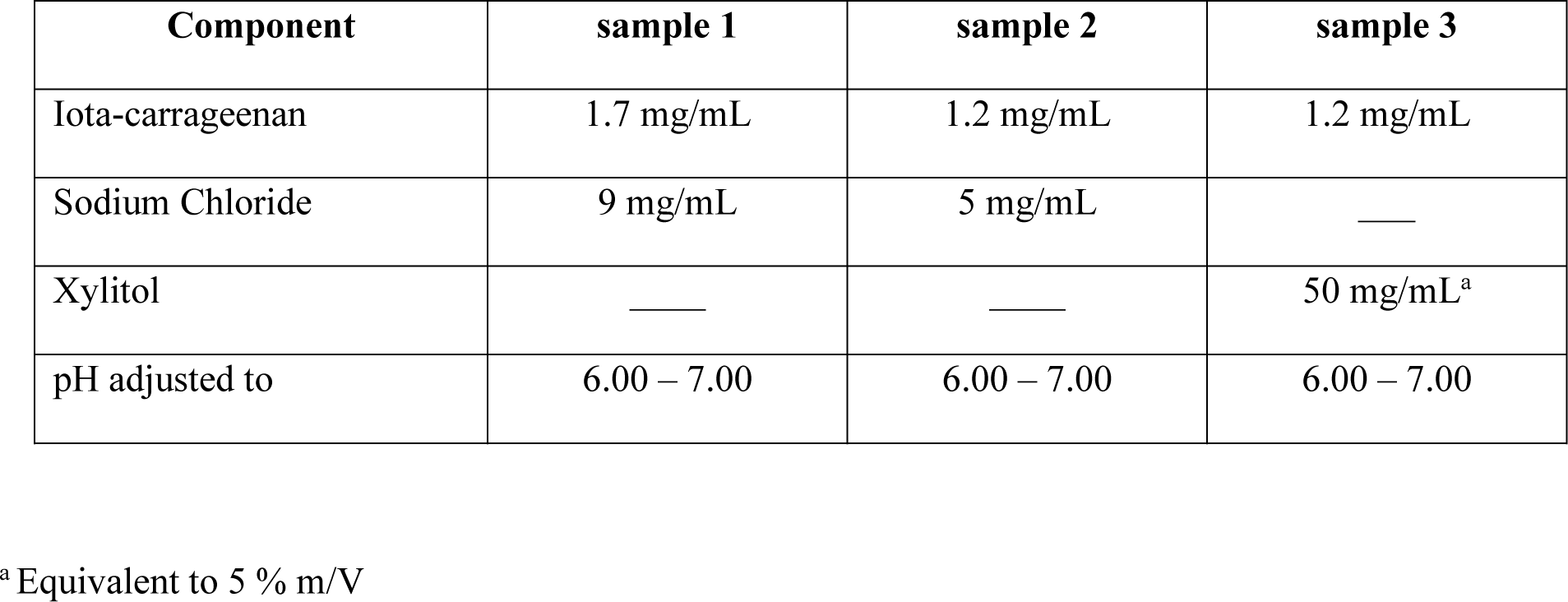
Composition of candidate nasal formulations (samples containing iota-carrageenan)

| Component | sample 1 | sample 2 | sample 3 |
| --- | --- | --- | --- |
| Iota-carrageenan | 1.7 mg/mL | 1.2 mg/mL | 1.2 mg/mL |
| Sodium Chloride | 9 mg/mL | 5 mg/mL | — |
| Xylitol | — | — | 50 mg/mL<sup>a</sup> |
| pH adjusted to | 6.00 – 7.00 | 6.00 – 7.00 | 6.00 – 7.00 |

<sup>a</sup> Equivalent to 5 % m/V

**Table 2.** Composition of placebo samples used as diluents (samples without iota-carrageenan)

| Component | sample P1 | sample P2 | sample P3 |
| --- | --- | --- | --- |
| Sodium Chloride | 9 mg/mL | 5 mg/mL | — |
| Xylitol | — | — | 50 mg/mL<sup>a</sup> |
| pH adjusted to | 6.00 – 7.00 | 6.00 – 7.00 | 6.0 – 7.00 |

<sup>a</sup> Equivalent to 5 % m/V

### Titer reduction assay

Vero E6 cells were seeded in 12-well plates at density of 2.5×10^5^/well and grown overnight at 370C under 5%CO2. Next day, cells were washed with PBS (pH 7.2), followed by addition of equivalent amount of c-MEM with reduced FBS (2%) and sample/placebo formulations. Formulations were incubated with cells for 2 hr, after which the supernatant was removed. Cells were infected with 2.5×10^4^ pfu (MOI=0.1) of virus for 1 hr at 37C, 5%CO2 with rocking at 15 min intervals. After incubation wells were washed with DPBS, and sample/placebo formulations were added at same concentrations. After incubation for 2 days, well contents were collected. For titer reduction, wells with no treatment (only virus) and cells only were included. Virus titer was determined by performing a TCID_50_ assay using MTT to measure cell viability. Virus endpoint titer was determined using the Reed-Muench formula and expressed as log TCID_50_/mL. Residual virus titer from sample/placebo formulation treated wells was plotted against virus titer from untreated wells.

## Results

To examine the antiviral effects of iota-carrageenan on SARS-CoV-2, three sample formulations were developed and tested. Each of the three sample formulations were tested in a dose dependent manner based on the concentration of iota-carrageenan and ranged from 600 µg/mL to 0 µg/mL. SARS-CoV-2 samples treated with 600 µg/mL and 60 µg/mL of sample formulation 1 were reduced > 3.75 Log when compared to untreated control (Figure 1). The 6 µg/mL concentration of sample formulation 1 also demonstrated an effect but to a lesser extent, with a 2.5 Log reduction in virus (Figure 1). No activity was observed with 0.6 µg/mL of Iota-carrageenan (Figure 1). Lastly, there was no reduction in virus with P1, suggesting that Iota-carrageenan and not the components of sample formulation 1 is inhibiting SARS-CoV-2 (Figure 1)

**Fig 1.**
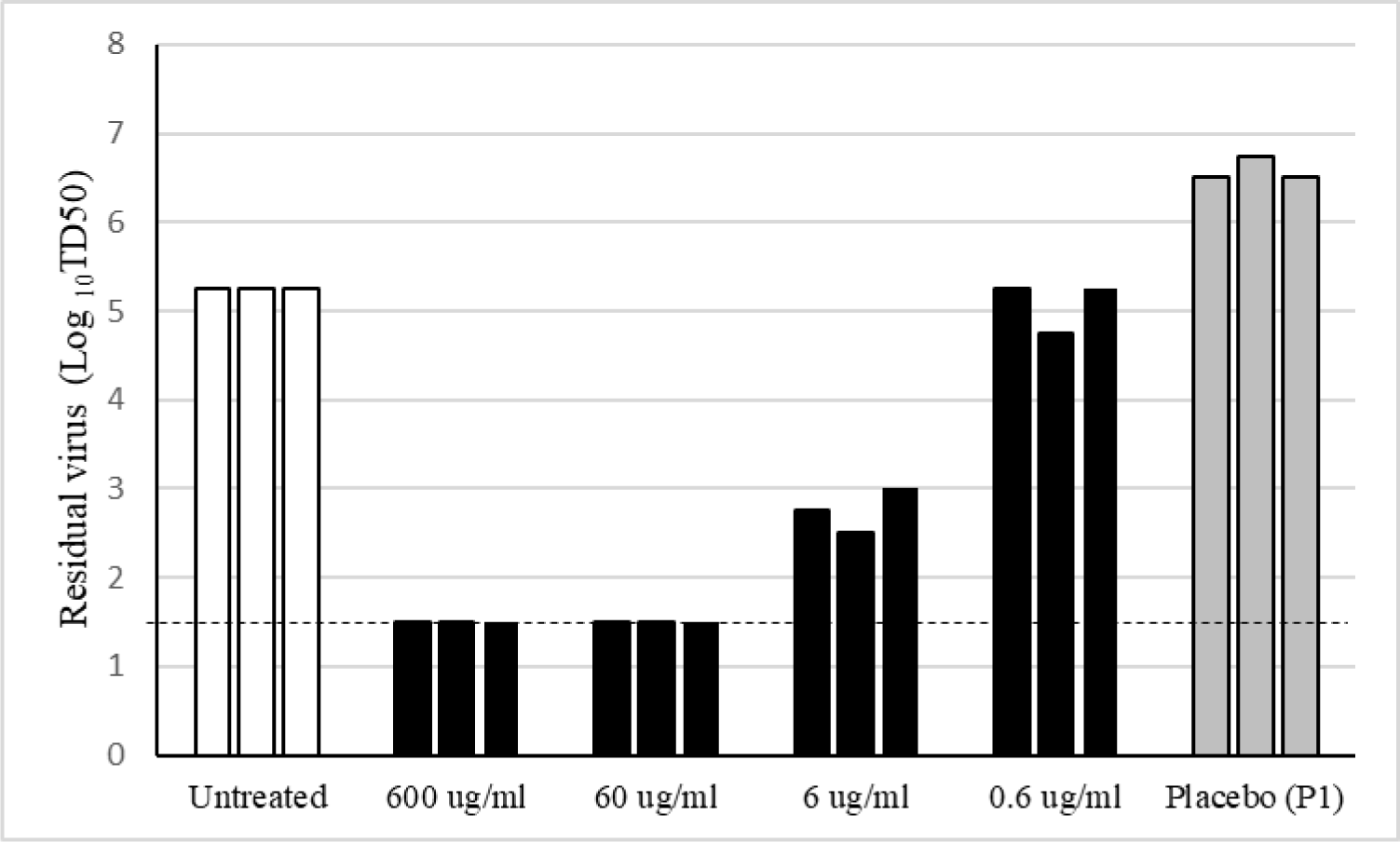
SARS-CoV-2 viral titer after treatment with samples 1 and P1 (3 replicates per treatment). Sample 1 composition: 1.7 mg/mL iota-carrageenan, 9 mg/mL sodium chloride, pH 6-7. Vero E6 were pre-treated with dilutions of sample 1 with sample P1 (placebo without iota-carrageenan) to get 600 µg/mL, 60 µg/mL, 6 µg/mL, 0.6 µg/mL iota-carrageenan final concentration for 2 h. After a 2 h pretreatment, cells were infected with SARS-CoV-2 and incubated for 48h in the presence of the same dilutions of sample 1. Supernatants were harvested and virus yield determined by an end point dilution assay (TCID50). Controls consisted of untreated infected cells or infected cells treated with P1 (no iota-carrageenan). Results were determined using the Reed and Muench formula and expressed as log TCID50/mL. Dotted line shows the limit of detection (LOD). Testing of samples was performed in triplicate. Underlying data reported in tables S3A and S3B as supporting information.

SARS-CoV-2 samples treated with dilutions of sample 2 at final iota-carrageenan concentrations of 600 µg/mL, 60 µg/mL, and 6 µg/mL were reduced > 4.25 Log compared to untreated control (Figure 2). The 0.6 µg/mL concentration of iota-carrageenan was also not effective with sample 2 (Figure 2). No reduction in virus with P2 was observed, suggesting that iota-carrageenan and not the components of formulation 2 is inhibiting SARS-CoV-2 (Figure 2).

**Fig 2.**
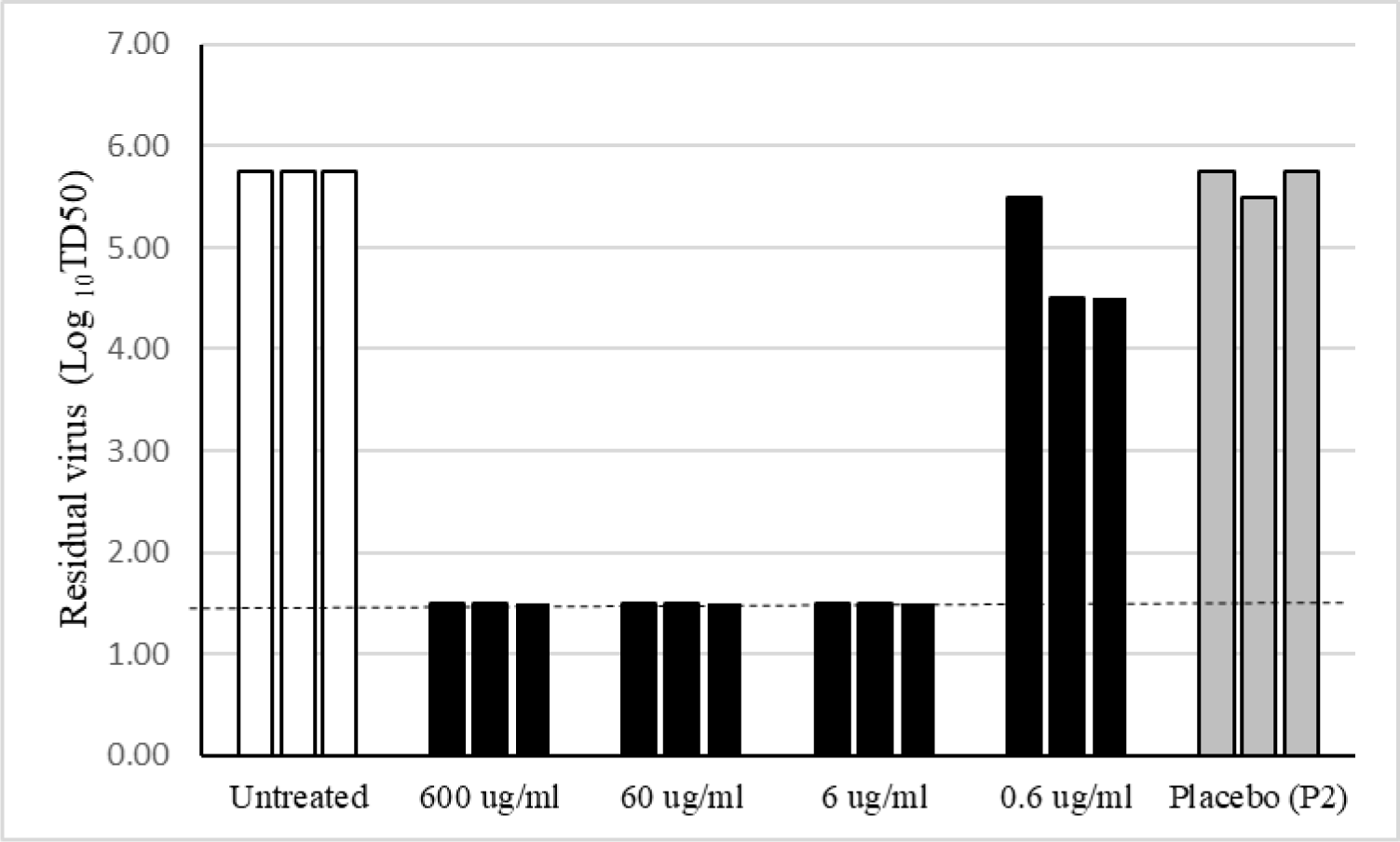
SARS-CoV-2 viral titer after treatment with samples 2 and P2 (3 replicates per treatment) Sample 2 composition: 1.2 mg/mL iota-carrageenan, 5 mg/mL sodium chloride, pH 6-7. Vero E6 were pre-treated with dilutions of sample 2 with sample P2 (placebo without iota-carrageenan) to get 600 μg/mL, 60 μg/mL, 6 μg/mL, and 0.6 μg/mL final iota-carrageenan concentration for 2 h. After a 2 h pretreatment, cells were infected with SARS-CoV-2 and incubated for 48h in the presence of the same dilutions of sample 2. Supernatants were harvested and virus yield determined by an end point dilution assay (TCID50). Controls consisted of untreated infected cells or infected cells treated with P2 (no iota-carrageenan). Results were determined using the Reed and Muench formula and expressed as log TCID50/mL. Dotted line shows the limit of detection (LOD). Testing of samples was performed in triplicate. Underlying data reported in tables S2A and S2B as supporting information.

SARS-CoV-2 samples treated with dilutions of sample 2 at final iota-carrageenan concentrations of 600 µg/mL, 60 µg/mL, and 6 µg/mL were reduced > 4.25 Log compared to untreated control (Figure 2). The 0.6 µg/mL concentration of iota-carrageenan was also not effective with sample 2 (Figure 2). No reduction in virus with P2 was observed, suggesting that iota-carrageenan and not the components of formulation 2 is inhibiting SARS-CoV-2 (Figure 2).

All concentrations tested (600 - 0.6 µg/mL) with sample 3 demonstrated antiviral activity including the P3 control that did not contain iota-carrageenan (Figure 3). Xylitol was present in this sample formulation and not in sample 1 or 2. The result suggests this component might also exert an antiviral effect.

**Fig 3.**
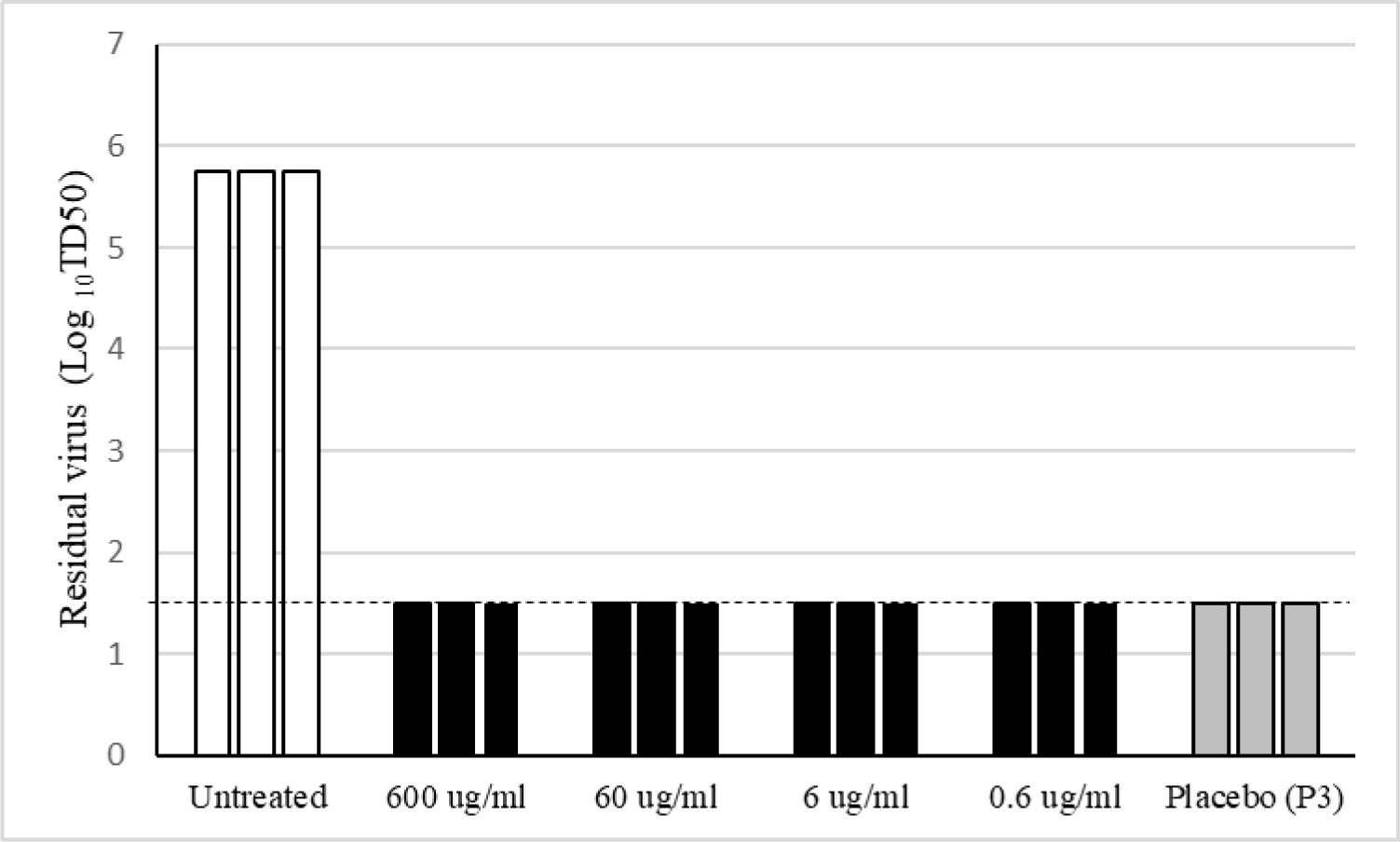
SARS-CoV-2 viral titer after treatment with samples 3 and P3 (3 replicates per treatment). Sample 3 composition: 1.2 mg/mL iota-carrageenan, 50 mg/mL xylitol, pH 6-7. Vero E6 were pre-treated with dilutions of sample 3 with sample P3 (placebo without iota-carrageenan) to get 600 μg/mL, 60 μg/mL, 6 μg/mL, and 0.6 μg/mL final iota-carrageenan concentration for 2 h. After a 2 h pretreatment, cells were infected with SARS-CoV-2 and incubated for 48h in the presence of the same dilutions of sample 3. Supernatants were harvested and virus yield determined by an end point dilution assay (TCID50). Controls consisted of untreated infected cells or infected cells treated with P3 (no iota-carrageenan). Results were determined using the Reed and Muench formula and expressed as log TCID50/mL. Dotted line shows the limit of detection (LOD). Testing of samples was performed in triplicate. Underlying data reported in tables S3A and S3B as supporting information.

A comparison of all three samples tested indicate that iota-carrageenan (600 µg/mL, 60 µg/mL, and 6 µg/mL) in samples 1 and 2 are effective at inhibiting SARS-CoV-2 (Table 3). Sample 3, which contained xylitol, was the most effective and demonstrated an antiviral effect at all concentrations tested (Table 3).

**Table 3.** Log reduction of TCID50/mL found after 48 hs.

| **Iota-carrageenan** | **Sample 1** | **Sample 2** | **Sample 3** |
| --- | --- | --- | --- |
| 600 µg/mL | ≥ 3.75 | ≥ 4.25 | ≥ 4.25 |
| 60 µg/mL | ≥ 3.75 | ≥ 4.25 | ≥ 4.25 |
| 6 µg/mL | 2.50 | ≥ 4.25 | ≥ 4.25 |
| 0.6 µg/mL | 0.17 | 0.92 | ≥ 4.25 |
| 0 µg/mL<sup>a</sup> | None | None | ≥ 4.25 |

The values reported are calculated as mean of untreated replicates minus mean of treated replicates. Mean values used for this calculation are reported in Tables S1B, S2B and S3B as supporting information. ^a^These are the placebo samples P1, P2 and P3 having the same components as samples 1, 2 and 3 except for iota-carrageenan.

## Discussion

Results from our study indicate that iota-carrageenan significantly inhibits SARS-CoV-2 in vitro. Our results are encouraging for clinical use of iota-carrageenan nasal spray for the prevention and early treatment of COVID-19. Clinical studies have demonstrated that iota-carrageenan nasal spray formulations already effective in vitro against rhinovirus [11] proved to be clinically effective in preventing and reducing symptoms and duration of common cold [14, 15, 16, 18]. Moreover, this study was designed so that the concentrations tested in vitro resemble the immediate concentration in nasal cavity and lower concentrations expected as iota-carrageenan is cleared from it. To do that we estimated airway surface liquid volume to be in the range 50 – 375 μL in nasal cavity based on a surface area of the nasal mucosa of 100 – 250 cm^2^ [28, 29, 30, 31] and airway surface liquid height estimated as 5 – 15 µm [32. 33]. If we take an average content of 200 μL of airway surface liquid in the nose plus 200 μL of formulation after delivering one 100-μL of a 1.2 mg/mL iota-carrageenan solution in each nostril, the immediate concentration of iota-carrageenan in the nasal cavity would be 600 µg/mL, coinciding with the highest concentration tested in vitro and capable of reducing virus yield to the LOD in our assay. Furthermore, considering that even 1/100 of this concentration is still active in vitro and that iota-carrageenan may stay for 4 hours [19] in the nasal cavity, there is a reasonable chance that this nasal spray may significantly help in the prevention and early treatment of COVID-19. Expected concentrations of iota-carrageenan in the nasal cavity will be even higher, if we consider a nasal formulation containing 1.7 mg / mL (0.17 % m/V) as some marketed nasal sprays.

The other remarkably interesting result is that xylitol exhibits antiviral activity on SARS-CoV-2 based on the results obtained with sample P3. Xylitol has been demonstrated to reduce titers of Human Respiratory Syncytial Virus in Hep-2 cells culture and in infected mice [24].

Despite the implementation of severe personal protection measures, pandemic continues to affect a significant proportion of health care workers with severe consequences for them, their patients, and the community. At the same time, most COVID-19 patients remain at home, thus increasing the likely exposure of household members and caregivers. Providing them with simple interventions as nasal sprays with either iota-carrageenan or xylitol in the same or different nasal devices may lower the risk of infection progression and transmission.

An inhalation solution of the same composition may be effective in case of severe cases of COVID 19. Even though there are studies showing safety of the use of both carrageenan and xylitol in nebulization [25,27], clinical trials would be needed to fully confirm these hypotheses. Risk of spreading the virus should be considered in this form of administration and due protection should be used to contain it [34].

We are starting multicenter randomized controlled trials to evaluate the efficacy of iota-carrageenan nasal sprays in health care staff assisting COVID-19 patients and in patients suffering from COVID 19 and other persons in close contact with them. However, it must be stressed that this and other similar nasal sprays are on the market and their safety profile is remarkable. The current COVID-19 emergency warrants the urgent development of potential strategies to protect people even if more robust data on antiviral therapies is yet to come.

## Conclusions

Iota-carrageenan inhibits SARS CoV-2 in vitro at concentrations easily achievable by nasal and nebulization formulations. Furthermore, xylitol exhibits antiviral activity on SARS-CoV-2, as well. An association with iota-carrageenan may be beneficial. There are already nasal sprays on the market having similar formulations to some of those tested in this in vitro study with adequate safety profile. Clinical trials are in progress to evaluate that nasal sprays based on the tested formulations are useful in the prevention and treatment of COVID 19. The data presented here are certainly encouraging in this direction.

## Acknowledgements

The authors thank Augusto Pich Otero and his team at Laboratorio Pablo Cassará S.R.L. (Argentina) for the supply of the sterile formulations for this in vitro study.

## References

1. Center for Systems Science and Engineering at Johns Hopkins University [Internet], COVID 19: Dashboard by the Center for Systems Science and Engineering at Johns Hopkins University. Available from: https://coronavirus.jhu.edu/map.html.

2. Wu Z, McGoogan J. Characteristics of and Important Lessons from the Coronavirus Disease 2019 (COVID-19) Outbreak in China: Summary of a Report of 72 314 Cases from the Chinese Center for Disease Control and Prevention. JAMA. 2020; 323(13): 1239–1242. doi:10.1001/jama.2020.2648.

3. Kelvin A, Halperin S. COVID-19 in children: the link in the transmission chain. The Lancet Infectious Diseases. 2020; 20: 633–634. doi: 10.1016/S1473-3099(20)30236-X.

4. Zou L, Ruan F, Huang M, Liang L, Huang H, Hong Z, et al. SARS-CoV-2 Viral Load in Upper Respiratory Specimens of Infected Patients. N Engl J Med. 2020; 382: 1177–1179. doi: 10.1056/NEJMc2001737.

5. Callahan C, Lee R, Lee G, Zulauf K, Kirby J, Arnaout R. Nasal-Swab Testing Misses Patients with Low SARS-CoV-2 Viral Loads. medRxiv [Preprint]. 2020. Available from: https://www.medrxiv.org/content/10.1101/2020.06.12.20128736v1. doi:10.1101/2020.06.12.20128736.

6. Liu Y, Liao W, Wan L, Xiang T, Zhang W. Correlation Between Relative Nasopharyngeal Virus RNA Load and Lymphocyte Count Disease Severity in Patients with COVID-19. Viral Immunology. 2020. doi: 10.1089/vim.2020.0062.

7. Liu Y, Yan L, Wan L, Xiang T, Le A, Liu J, et al. Viral dynamics in mild and severe cases of COVID-19. The Lancet Infectious Diseases. 2020; 20: 656–657. doi: 10.1016/S1473-3099(20)30232-2.

8. Ahmadi A, Zorofchian S, Abu Bakar S, Zandi K. Antiviral Potential of Algae Polysaccharides Isolated from Marine Sources: A Review. BioMed Research International. 2015. doi: 10.1155/2015/825203

9. Buck CB, Thompson CD, Roberts JN, Müller M, Lowry DR, et al. Carrageenan is a potent inhibitor of papillomavirus infection. PLoS Pathog. 2006; 2 (7): e69. doi: 10.1371/journal.ppat.0020069.

10. Girond S, Crance JM, Van Cuyck-Gandre H, Renaudet J, Deloince R. Antiviral activity of carrageenan on hepatitis A virus replication in cell culture. Res Virol 1991; 142 (4): 261–270. doi:10.1016/0923-2516(91)90011-q.

11. Grassauer A, Weinmuellner R, Meier C, Pretsch A, Prieschl-Grassauer E, Unger H. lota-Carrageenan is a potent inhibitor of rhinovirus infection. Virol Journal 2008; 5:107. doi:10.1186/1743-422X-5-107.

12. Shao Q, Guo Q, Xu Wp, Li Z, Zhao Tt, Specific inhibitory effect of κ-carrageenan polysaccharide on swine pandemic 2009 H1N1 influenza virus. PLoS One. 2015; 10 (5): e0126577. doi: 10.1371/journal.pone.0126577.

13. Talarico LB, Damonte EB. Interference in dengue virus adsorption and uncoating by carrageenans. Virology. 2007; 363: 473–485. doi: 10.1016/j.virol.2007.01.043.

14. Eccles R, Meier C, Jawad M, Weinmüller R, Grassauer A, Prieschl-Grassauer E. Efficacy and safety of an antiviral Iota-Carrageenan nasal spray: A randomized, double-blind, placebo-controlled exploratory study in volunteers with early symptoms of the common cold. Respiratory research 2010: 11. 108. 10.1186/1465-9921-11-108.

15. Koenighofer M, Lion T, Bodenteich A, Prieschl-Grassauer E, Grassauer A, Unger H et al. Carrageenan nasal spray in virus confirmed common cold: Individual patient data analysis of two randomized controlled trials. Multidisciplinary Respiratory Medicine. 2014; 9: 57. doi:10.1186/2049-6958-9-57.

16. Ludwig M, Enzenhofer E, Schneider S, Rauch M, Bodenteich A, Neumann K, et al. Efficacy of a Carrageenan nasal spray in patients with common cold: A randomized controlled trial. Respiratory Research. 2013; 14: 124. doi:10.1186/1465-9921-14-124.

17. Leibbrandt A., Meier C, König-Schuster M, Weinmüller R, Kalthoff D, Pflugfelder B, et al. Iota-Carrageenan Is a Potent Inhibitor of Influenza A Virus Infection, PLoS ONE. 2010; 5(12): e14320. doi: 10.1371/journal.pone.0014320.

18. Fazekas T, Eickhoff Ph, Pruckner N, Vollnhofer G, Fischmeister G, Diakos C, et al. Lessons learned from a double-blind randomized placebo-controlled study with a iota-carrageenan nasal spray as medical device in children with acute symptoms of common cold. BMC Complementary and Alternative Medicine 2012; 12: 147. doi:10.1186/1472-6882-12-147.

19. Graf C, Bernkop-Schnürch A; Egyed A, Koller C, Prieschl-Grassauer E, Morokutti-kurz M. Development of a nasal spray containing xylometazoline hydrochloride and iota-carrageenan for the symptomatic relief of nasal congestion caused by rhinitis and sinusitis. International Journal of General Medicine. 2018; 11: 275–283. doi: 10.2147/IJGM.S167123.

20. Morokutti-Kurz M; König-Schuster M, Koller C, Graf C, Graf Ph, Kirchoff N, et al. The Intranasal Application of Zanamivir and Carrageenan Is Synergistically Active against Influenza A Virus in the Murine Model. PLoS ONE. 2015; 10(6): e0128794. doi: 10.1371/journal.pone.0128794.

21. Salli K, Lehtinen M, Tiihonen K, Ouwehand A. Xylitol’s Health Benefits beyond Dental Health: A Comprehensive Review. Nutrients. 2019; 11: 1813. doi: 10.3390/nu11081813

22. Sakallioğlu, Ö, Güvenç A, Cingi C. Xylitol and its usage in ENT practice. The Journal of Laryngology & Otology. 2014; 128: 580–585. doi: 10.1017/S0022215114001340.

23. Lin L, Tang X, Wei J, Dai F, Sun G (2017). Xylitol nasal irrigation in the treatment of chronic rhinosinusitis. American journal of otolaryngology. 38. 10.1016/j.amjoto.2017.03.006.

24. Xu ML, Wi GM, Kim HJ, Kim HJ. Ameliorating Effect of Dietary Xylitol on Human Respiratory Syncytial Virus (hRSV) Infection, Biol. Pharm. Bull. 2016; 39: 540–546. doi:10.1248/bpb.b15-00773.

25. Hebar A, Koller C, Seifert JM, Chavicovsky M, Bodenteich A, Bernkop Schnürch A, et al. Non-Clinical Safety Evaluation of Intranasal Iota-Carrageenan. PLOS ONE. 2015; 10 (4): e0122911. 10.1371/journal.pone.0122911.

26. Weissman JD, Fernandez F, Hwang PH. Xylitol Nasal Irrigation in the Management of Chronic Rhinosinusitis: A Pilot Study, Laryngoscope. 2011; 121:2468–2472. doi: 10.1002/lary.22176.

27. Durairaj L, Launspach, J, Watt JL, Busing TR, Kline JN, Thorne PS, et al. Safety assessment of inhaled xylitol in mice and healthy volunteers. Respiratory Research. 2004; 5: 13. doi: 10.1186/1465-9921-5-13.

28. Bitter C, Suter-Zimmermann K, Surber C. Nasal Drug Delivery in Humans, in Surber C, Elsner P, Farage MA, editors. Topical Applications and the Mucosa. Curr Probl Dermatol. Basel, Karger, 2011; 40: 20–35. doi:10.1159/000321044.

29. Garcia GJM, Schroeter JD, Segal RA, Stanek J, Foureman GL, Kimbell JS. Dosimetry of nasal uptake of water-soluble and reactive gases: A first study of interhuman variability, Inhalation Toxicology, 2009; 21(7): 607–618, doi: 10.1080/08958370802320186

30. Gizurarzon S. Anatomical and Histological Factors Affecting Intranasal Drug and Vaccine Delivery, Current Drug Delivery. 2012; 9: 566–582. doi: 10.2174/156720112803529828.

31. Pires A, Fortuna A, Alves G, Falcão A. Intranasal Drug Delivery: How, Why and What for?, J Pharm Pharmaceut Sci. 2009; 12(3): 288–311. doi: 10.18433/j3nc79

32. Helassa N, Garnett JP, Farrant M, Khan F, Pickup JC, Hahn KM, et al. A novel fluorescent sensor protein for detecting changes in airway surface liquid glucose concentration Biochem. J. 2014; 464: 213–220, doi:10.1042/BJ20141041.

33. Wagenman M, Naclerio RM (1992), Anatomic and physiologic considerations in sinusitis, J Allergy Clin Immunol, 1992; 90; 3 (Part 2): 419–423

34. Ari A. Practical strategies for a safe and effective delivery of aerosolized medications to patients with COVID-19. Respiratory Medicine. 2020;167: 105987. doi: 10.1016/j.rmed.2020.105987.

