## Supplementary material for "Iota-carrageenan and Xylitol inhibit SARS-CoV-2 in cell culture": Tables S

### Supporting information

**Table S1A. Original data for Fig 1**

| Treatment | TCID50/mL | | |
|---|---|---|---|
| | **Replicate 1** | **Replicate 2** | **Replicate 3** |
| Untreated | 1.78E+05 | 1.78E+05 | 1.78E+05 |
| 600 µg/ml | 3.16E+01 | 3.16E+01 | 3.16E+01 |
| 60 µg/ml | 3.16E+01 | 3.16E+01 | 3.16E+01 |
| 6 µg/ml | 5.62E+02 | 3.16E+02 | 1.00E+03 |
| 0.6 µg/ml | 1.78E+05 | 5.62E+04 | 1.78E+05 |
| Placebo (P1) | 3.16E+06 | 5.62E+06 | 3.16E+06 |

**Table S1B. Decimal logarithm of values of TCID50 used to build Fig 1 and Table 3**

| Treatment | $LOG_{10}$TCID50/mL | | | |
|---|---|---|---|---|
| | **Replicate 1** | **Replicate 2** | **Replicate 3** | **Mean** |
| Untreated | 5.25 | 5.25 | 5.25 | 5.25 |
| 600 µg/ml | 1.50 | 1.50 | 1.50 | 1.50 |
| 60 µg/ml | 1.50 | 1.50 | 1.50 | 1.50 |
| 6 µg/ml | 2.75 | 2.50 | 3.00 | 2.75 |
| 0.6 µg/ml | 5.25 | 4.75 | 5.25 | 5.08 |
| Placebo (P1) | 6.50 | 6.75 | 6.50 | 6.58 |

**Table S2A. Original data for Fig 2**

| Treatment | TCID50/mL | | |
|---|---|---|---|
| | **Replicate 1** | **Replicate 2** | **Replicate 3** |
| Untreated | 5.62E+05 | 5.62E+05 | 5.62E+05 |
| 600 µg/ml | 3.16E+01 | 3.16E+01 | 3.16E+01 |
| 60 µg/ml | 3.16E+01 | 3.16E+01 | 3.16E+01 |
| 6 µg/ml | 3.16E+01 | 3.16E+01 | 3.16E+01 |
| 0.6 µg/ml | 3.16E+05 | 3.16E+04 | 3.16E+04 |
| Placebo (P2) | 5.62E+05 | 3.16E+05 | 5.62E+05 |

**Table S2B. Decimal logarithm of values of TCID50 used to build Fig 2 and Table 3**

| Treatment | LOG$_{10}$TCID50/mL | | | |
|---|---|---|---|---|
| | **Replicate 1** | **Replicate 2** | **Replicate 3** | **Mean** |
| Untreated | 5.75 | 5.75 | 5.75 | 5.75 |
| 600 µg/ml | 1.50 | 1.50 | 1.50 | 1.50 |
| 60 µg/ml | 1.50 | 1.50 | 1.50 | 1.50 |
| 6 µg/ml | 1.50 | 1.50 | 1.50 | 1.50 |
| 0.6 µg/ml | 5.50 | 4.50 | 4.50 | 4.83 |
| Placebo (P2) | 5.75 | 5.50 | 5.75 | 5.67 |

**Table S3A. Original data for Fig 3**

| Treatment | TCID50/mL | | |
|---|---|---|---|
| | **Replicate 1** | **Replicate 2** | **Replicate 3** |
| Untreated | 5.62E+05 | 5.62E+05 | 5.62E+05 |
| 600 µg/ml | 3.16E+01 | 3.16E+01 | 3.16E+01 |
| 60 µg/ml | 3.16E+01 | 3.16E+01 | 3.16E+01 |
| 6 µg/ml | 3.16E+01 | 3.16E+01 | 3.16E+01 |
| 0.6 µg/ml | 3.16E+01 | 3.16E+01 | 3.16E+01 |
| Placebo (P3) | 3.16E+01 | 3.16E+01 | 3.16E+01 |

**Table S3B. Decimal logarithm of values of TCID50 used to build Fig 3 and Table 3**

| Treatment | LOG$_{10}$TCID50/mL | | | |
|---|---|---|---|---|
| | **Replicate 1** | **Replicate 2** | **Replicate 3** | **Mean** |
| Untreated | 5.75 | 5.75 | 5.75 | 5.75 |
| 600 µg/ml | 1.50 | 1.50 | 1.50 | 1.50 |
| 60 µg/ml | 1.50 | 1.50 | 1.50 | 1.50 |
| 6 µg/ml | 1.50 | 1.50 | 1.50 | 1.50 |
| 0.6 µg/ml | 1.50 | 1.50 | 1.50 | 1.50 |
| Placebo (P3) | 1.50 | 1.50 | 1.50 | 1.50 |
